# Exploring multi-level microbial interactions from individual 3D genomes to community networks

**DOI:** 10.64898/2025.12.03.692222

**Authors:** Wen-Jie Jiang, Kang-Wen Cai, Yuan-Chen Sun, Zhetao Zheng, Fangdi Xu, Rui-Xiang Gao, NaNa Wei, Hanwen Zhu, Yu-Juan Wang, Qing Xia, Chang Lu, Ming Xu, Hua-Jun Wu

## Abstract

Microbial communities interact with their hosts through complex genomic networks that influence ecosystem stability and disease progression. Here, we present FindMeta3D, a computational framework that simultaneously identifies microbial three-dimensional (3D) genome structures and cross-domain interaction networks. Applying this approach to 528 Hi-C samples, we resolved 3D genome structures of 344 microbial species, revealing five evolutionarily conserved chromatin folding patterns linked to intrinsic sequence features. Further analyses demonstrated distinct microbial-host interaction preferences and identified functional interaction hotspots that are critical for infection. Experimental deletion of such hotspots in the EBV genome resulted in significant infection defects, demonstrating their essential role in viral infectivity. Additionally, we constructed the first Cross-domain Microbial Interaction Network (CMIN), which uncovered pathogen-specific subnetworks and demonstrate dramatic restructuring of gut microbial communities in neutropenic patients, including enhanced *Klebsiella*-phage interactions. Subnetwork analysis identified potential phage therapy targets, such as *Klebsiella phage ST16-OXA48phi5.4*. These findings provide fundamental insights into microbial 3D genomics and establish FindMeta3D as a powerful platform for studying microbial genome structure and communities and developing antimicrobial strategies.

## Introduction

While significant advances have been made in mammalian 3D genome research^1–3^, the architecture of microbial genomes remains poorly understood. To date, only about 20 microbial species have had their 3D genomes characterized, severely limiting our knowledge of microbial chromatin organization. Microbial communities engage in complex interactions with their hosts that are central to maintaining ecosystem balance and modulating disease progression^4–7^. Inter-species genomic contacts—including direct DNA-DNA interactions—serve as key mechanisms coordinating metabolic cooperation, resource competition, and niche partitioning^8–10^. Current methods, such as culture-based approaches, can only examine specific microbial pairs^11^, while metagenomic sequencing relies on indirect inference, not capturing physical interaction events directly^12^. Although archaea, bacteria, and fungi are known to influence hosts through DNA-mediated interactions^13,14^, systematic tools to study these processes are lacking. While Hi-C technology enables genome-wide interaction profiling^15^, existing studies focus predominantly on individual virus-host and phage-bacteria interactions^16–18^, leaving most microbial interaction networks unexplored.

To address these gaps, we develop FindMeta3D, an analytical framework that simultaneously resolves microbial intra-/inter-species interactions and host-microbe contacts through rigorous quality control and iterative alignment strategies. Applying FindMeta3D to 528 Hi-C samples, we: 1) resolved high-quality 3D genomes for 344 microbial species that form five distinct structural types, associating with intrinsic sequence properties; 2) characterized the contacting preferences of different microbial species to specific human genomic regions; 3) identified microbial genomic hotspots interacting with human genome, and validate by deletion experiments showing significantly reduced infectivity; and 4) constructed the first Cross-domain Microbial Interaction Network (CMIN) spanning archaea, bacteria, fungi, and viruses, revealing 25 subnetworks involving WHO-listed pathogens. Notably, comparing gut microbial networks between healthy and neutropenic individuals uncovered enhanced *Klebsiella*-phage interactions during infection, nominating potential therapeutic phages (e.g., *Klebsiella phage ST16-OXA48phi5.4*). Collectively, FindMeta3D establishes a scalable platform for microbial interactome studies and highlights the regulatory importance of cross-domain interaction networks in microbial communities.

## Results

### Overview of FindMeta3D

Here, we developed FindMeta3D, an algorithm specifically designed to systematically mine 3D genome folding of individual microbes, and microbial-microbial and microbial-host interactions through Hi-C data (Figure 1a). The FindMeta3D workflow comprises three critical steps: First, we performed rigorous quality control on raw Hi-C data to filter out low-quality reads. Next, the high-quality paired-end reads were aligned to the telomere-to-telomere human reference genome (CHM13, T2T assembly)^19^, followed by removal of reads where both ends map to the host genome (representing host-host interactions). Finally, to precisely resolve interaction types, the remaining reads were iteratively aligned to reference genomic databases covering four major microbial domains—Archaea, Bacteria, Fungi, and Viruses. We additionally performed multi-mapped reads detection to eliminate sequences that map to multiple species, thereby effectively removing cross-species contamination signals. FindMeta3D provides a robust computational framework for elucidating complex interaction networks within microbial communities and between microbes and their host genomes.

**Figure 1.**
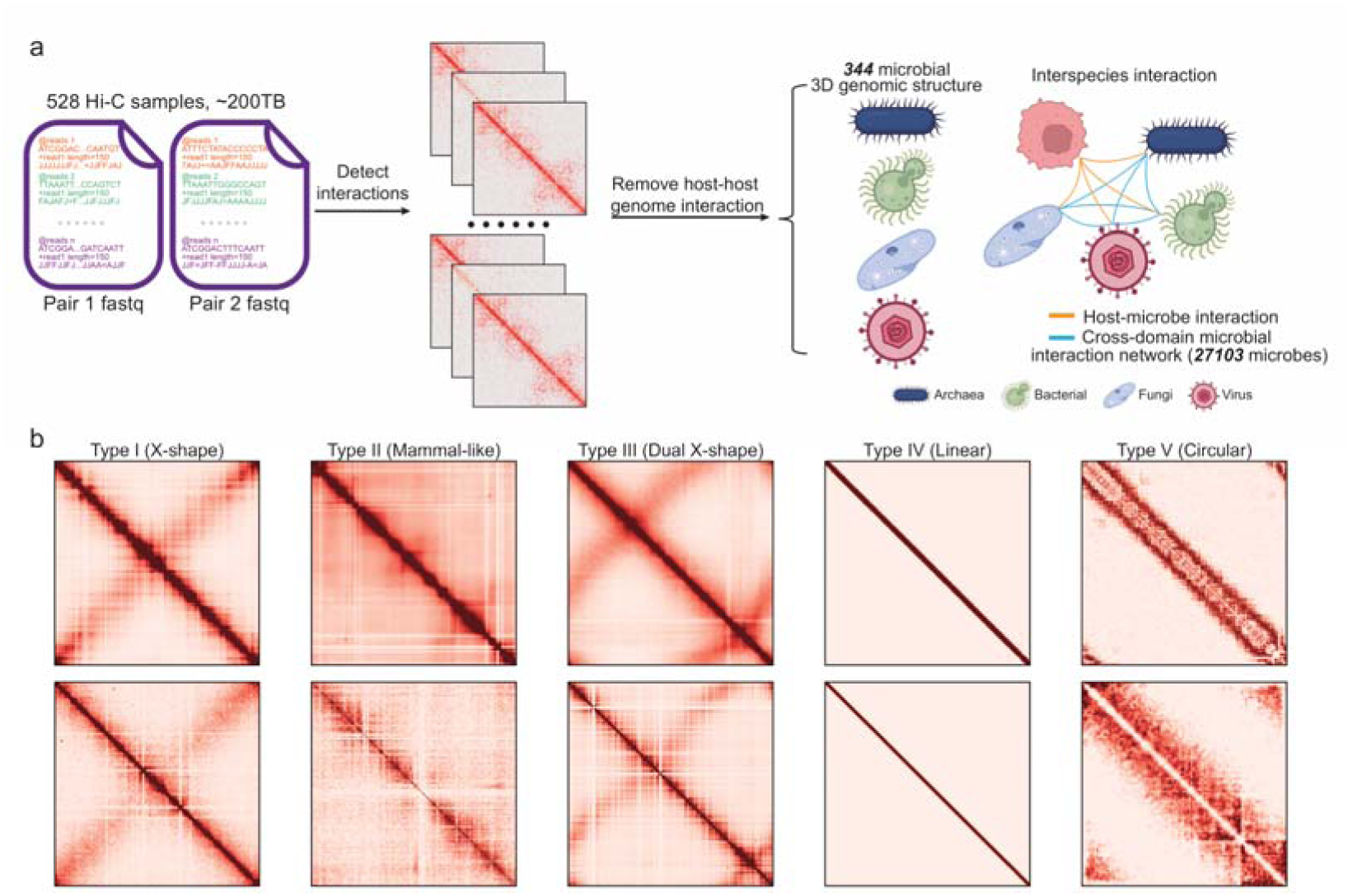
Computational framework of FindMeta3D and classification of microbial 3D genome architectures. (a) Schematic overview of FindMeta3D. (b) Five distinct patterns of microbial 3D genome structure: Type I (X-shape), Type II (Mammal-like), Type III (Dual X-shape), Type IV (Linear), and Type V (Circular).

### Five Classes of Microbial 3D Genome Structures

By analyzing 528 Hi-C samples using FindMeta3D, we resolved 3D genome structures of 344 microbial species, revealing five characteristic nuclear architecture patterns at the genome scale: Type I (X-shape), Type II (Mammal-like), Type III (Dual X-shape), Type IV (Linear), and Type V (Circular) (Figure 1b, Figure S1a). To better understand these folding principles, we performed 3D modeling^20^ on representative species from each architectural type (Figure S1a). Our modeling results demonstrated clear organizational differences among these structural classes. Types I-III exhibited highly condensed architectures characterized by extensive distal chromatin interactions. (Figure 1b). In contrast, Types IV and V displayed more open conformations with distinct organizational features. Type IV structures showed predominant local interactions with minimal long-range contacts, while Type V formed complete circular topologies maintained by robust terminal chromatin interactions (Figure 1b). These structural variations likely reflect fundamental differences in genome packaging strategies across microbial species.

To account for potential biases introduced by variations in genome size during microbial 3D genome classification, we implemented a standardized analytical pipeline. First, we normalized all 344 microbial interaction matrices to a uniform 256 × 256 resolution using an interpolation algorithm. We then performed feature extraction using linear discriminant analysis (LDA)^21^ followed by dimensionality reduction and visualization with UMAP (Figure 2a,b). This approach revealed a clear separation of the five structural types in low-dimensional space, confirming their distinct organizational patterns. We systematically examined the association between 3D genome architectures and fundamental genomic characteristics by analyzing three key parameters: GC content, genome length, and DNA bendability^22^ (Figure 2c-e). Our analysis identified significant differences across structural types: Type I and III genomes possessed both the highest GC content and longest genomic lengths, while Type V genomes showed the opposite trend with the lowest GC content and shortest lengths. Notably, Type V genomes exhibited markedly elevated DNA bendability compared to other types, suggesting their circular conformation may be stabilized through enhanced local DNA flexibility. These findings establish unprecedented associations between 3D genome organization and intrinsic sequence properties across microbial species.

**Figure 2.**
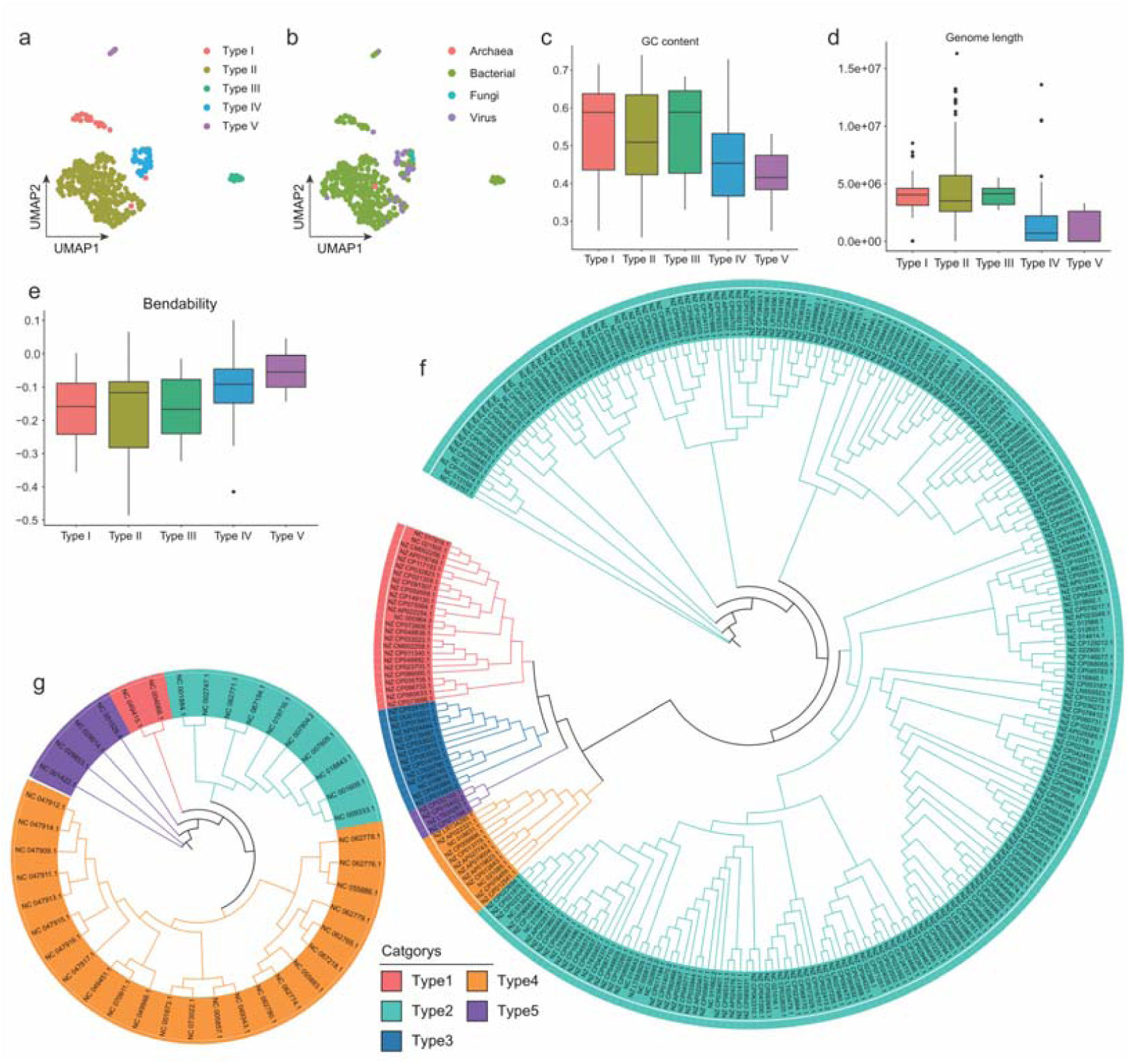
Genomic and evolutionary features linked to microbial 3D genome architecture. (a) UMAP projection showing distinct clustering of the five structural types (Type I-V) across 344 microbial genomes. (b) Corresponding taxonomic distribution of microbial species. (c-e) Comparative analysis of three intrinsic genomic features across structural types: (c) GC content (%), (d) genome length (bp), and (e) DNA bendability. Boxplots represent median values with interquartile ranges. (f-g) Phylogenetic distribution of structural types in (f) bacterial and (g) viral, demonstrating evolutionary conservation of 3D genome organization.

### Evolutionary Conservation of 3D Genome Structures Across Microbial Species

The structural maintenance of chromosomes (SMC) and condensin protein complexes are known to play pivotal roles in organizing chromosomal architectures^23–25^. To examine the evolutionary relationships underlying structural conservation, we conducted comprehensive multiple sequence alignments of SMC and condensin coding regions across all 301 bacterial genomes. Using conserved domain features, we reconstructed a robust maximum likelihood phylogenetic tree with IQ-TREE^26^ (Figure 2f). Our evolutionary analysis demonstrated striking phylogenetic clustering of bacteria sharing identical 3D genome structures (bootstrap value >95%), while distinct structural types showed clear evolutionary divergence. Notably, phylogenetic reconstruction revealed close relationships between the two X-shaped structural groups, displaying the greatest genetic divergence from mammal-like structural types. Phylogenetic analysis of 41 viral genomes similarly revealed closer evolutionary distances among viruses sharing identical 3D genome structures (Figure 2g), reinforcing the general principle that 3D genome organization reflects phylogenetic relationships across diverse microbial lineages.

### ACP contributes to Bacterial 3D Genome Architecture

To identify potential protein determinants of bacterial 3D genome organization, we systematically analyzed 301 bacterial genomes representing all five structural types. While SMC and condensin proteins are known chromatin regulators, our BLAST analysis revealed a weak association between their presence in a species and its structural types (Figure S2a). This suggests these canonical structural proteins alone cannot fully explain the observed architectural diversity. Intriguingly, acyl carrier protein (ACP) exhibited the most diverse distribution across structural types, with a limited subset of Type IV species retaining detectable ACP orthologs (Figure S2b). For those Type IV species possessing ACP, multiple sequence alignment analysis uncovered distinct amino acid substitutions not observed in other structural types (Figure S2c). Structural modeling of these Type IV-specific ACP variants predicted unique conformational rearrangements (Figure S2d,e) that may impair long-range chromatin interactions, potentially explaining the characteristic linear architecture of Type IV genomes.

### Microbial Targeting of Specific Chromatin in Human Genome

Using microbial-human genome interaction maps identified by FindMeta3D, we systematically quantified the interaction frequency of diverse microbes to the human genome. Viral genomes demonstrated the highest interaction frequency among all microbial groups (Figure 3a). Integration of multiple epigenetic datasets, including H3K36me3, H3K4me3, ATAC-seq, H3K27ac, H3K9me3, replication timing, nucleolar-associated domains (NADs), common fragile sites (CFSs), and lamina-associated domains (LADs), revealed distinct patterns of microbial-interacting human genome preferences. Most microbes preferentially interact with genomic regions enriched by H3K36me3, H3K4me3, H3K27ac, open chromatin and early replication timing, while avoiding H3K9me3 regions (Figure 3b). However, approximately 15% of microbes, primarily fungi and archaea, exhibited an opposite pattern - showing an interacting preference for inactive chromatin regions and NADs/LADs (Figure 3b). These unique interaction preferences may represent an evolutionary adaptation allowing these microbes to establish specialized niches within different chromatin regions of the host nucleus.

**Figure 3.**
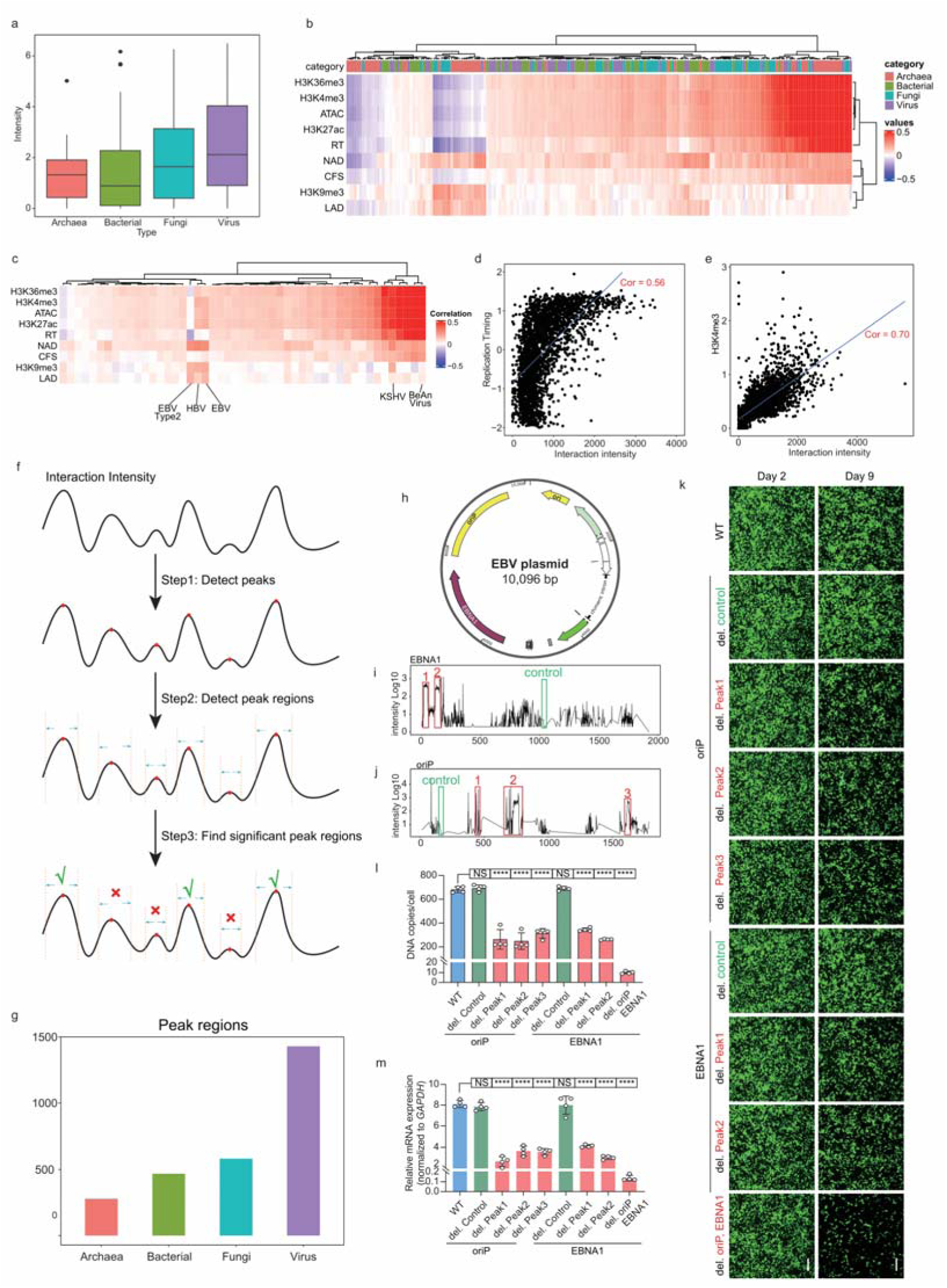
Characterization and functional validation of microbial-human genomic interactions. (a) Normalized interaction frequency between the genomes of four microbial domains (Archaea, Bacteria, Fungi, and Viruses) and human. (b, c) Genome-wide correlation heatmap between microbial (b) or viral (c) interaction frequency (y-axis) and nine human epigenetic features (x-axis) calculated in 1-Mb genomic windows. (d,e) Scatter plots showing the correlation between *BeAn* viral interaction frequency and (d) replication timing or (e) H3K4me3 signals. (f) Schematic workflow of the FindMetaPeaks algorithm. (g) Distribution of 2,757 significant interaction hotspots across microbial domains. (h) Schematic of EBV-derived vector. (i, j) Significant interaction hotspots and control regions in EBV-derived vector. (k) Fluorescence microscopy of HEK293T cells on day 2 and 9 after transfection with various EBV-derived vectors. (l, m) Quantitative analysis of (l) DNA copy number (qPCR) and (m) relative mRNA expression (RT-qPCR) on day 9 after transfection (\*\*\*\**p* < 0.0001, NS: not significant).

Viruses, as the most prevalent microbial group infecting human cells, displayed a particularly strong preference for active chromatin regions (Figure 3c). Most viral genomes preferentially interact with transcriptionally active, open chromatin regions. For instance, the *BeAn virus* showed particularly strong correlations with active chromatin marks: ATAC (r=0.68), H3K27ac (r=0.58), H3K36me3 (r=0.70), replication timing (r=0.56), and H3K4me3 (r=0.70) (Figure 3d,e; Figure S3a-c). These striking correlations suggest that the *BeAn virus* may have evolved to specifically target transcriptionally active, early-replicating, and open chromatin regions of the host genome - a strategy that likely facilitates efficient viral gene expression while evading host immune surveillance.

### Identification of High-Frequency Interacting hotspots in Microbial Genomes

Further analysis revealed that microbial-human genome interactions exhibited non-random distribution patterns also on the microbe side, with specific genomic loci demonstrating preferential interaction frequencies. To systematically map these critical interaction hotspots in microbial genomes, we developed FindMetaPeaks, a computational algorithm that identifies hotspots based on interaction intensity distributions through False Discovery Rate (FDR) correction (q < 0.05) and a minimum interaction intensity threshold (Z-score > 2) (Figure 3f). Application of this algorithm across archaeal, bacterial, fungal, and viral genomes identified 2,757 significant interaction hotspots, with viral genomes contributing the majority (1,429 regions, 51.8%) of these hotspots (Figure 3g).

To investigate the effect of these interaction hotspots on virus infection, we employed an oriP/EBNA1 vector derived from Epstein-Barr virus (EBV) as our experimental model. The vector enabled stable expression of green fluorescent protein (GFP) in mammalian cells, which depends on the continuous extrachromosomal replication requiring the cis-acting oriP element and the trans-acting EBNA1 gene (Figure 3h). FindMetaPeaks identified two significant interaction hotspots within the EBNA1 gene region (Figure 3i) and three in the oriP gene region (Figure 3j). We generated mutant EBV-derived vectors with individual hotspot deletions, and compared them to that with control regions and complete gene deletions. While initial transfection efficiencies (day 2) were comparable across all groups, by day 9 after transfection we observed significant phenotypic differences: vectors lacking hotspots showed markedly reduced fluorescence (Figure 3k), accompanied by a 50.4% decrease in DNA replication (qPCR, *p* < 0.0001) (Figure 3l) and 60% reduction in mRNA expression (RT-qPCR, *p* < 0.0001) (Figure 3m). In contrast, control region deletions showed no significant effects (*p* = 0.9994), suggesting the potential contribution of interaction hotspots to maintain DNA replication during transgene expression of viral vectors. These results demonstrate that interaction hotspots in viral genome may play essential roles in sustaining long-term infection through regulation of DNA replication and gene expression, highlighting their critical importance in maintaining persistent host-pathogen interactions.

### The First Cross-Domain Microbial Interaction Network

Through systematic application of the FindMeta3D algorithm on the collected 528 Hi-C samples, we extracted and statistically tested all inter-species interactions across microbial domains, enabling the construction of the first Cross-domain Microbial Interaction Network (CMIN) (Figure 4a). The network comprised 27,103 nodes spanning Archaea, Bacteria, Fungi, and Viruses, connected by 68,624 statistically significant interaction edges. Network topology analysis identified *Aeribacillus phage AP45* as a central hub exhibiting remarkable connectivity, forming interactions with 728 archaeal, 1,070 bacterial, 884 fungal, and 172 viral genomes, achieving a degree centrality of 2,854 (Figure S4a). At the other extreme, we observed highly specific interactions demonstrating remarkable partner selectivity, including the exclusive pairing between *Bacteroides intestinalis* and *Bacteroides phage B124-14*, and the unique interaction between *Lactococcus phage P596* and *Lactococcus lactis subsp. Lactis* (Figure S4b,c). These specific interactions likely reflect evolutionarily conserved microbial partnerships. The CMIN provides not only a comprehensive map of microbial interactomes but also a transformative framework for identifying potential synergistic pathogenic factors through systematic network analysis.

**Figure 4.**
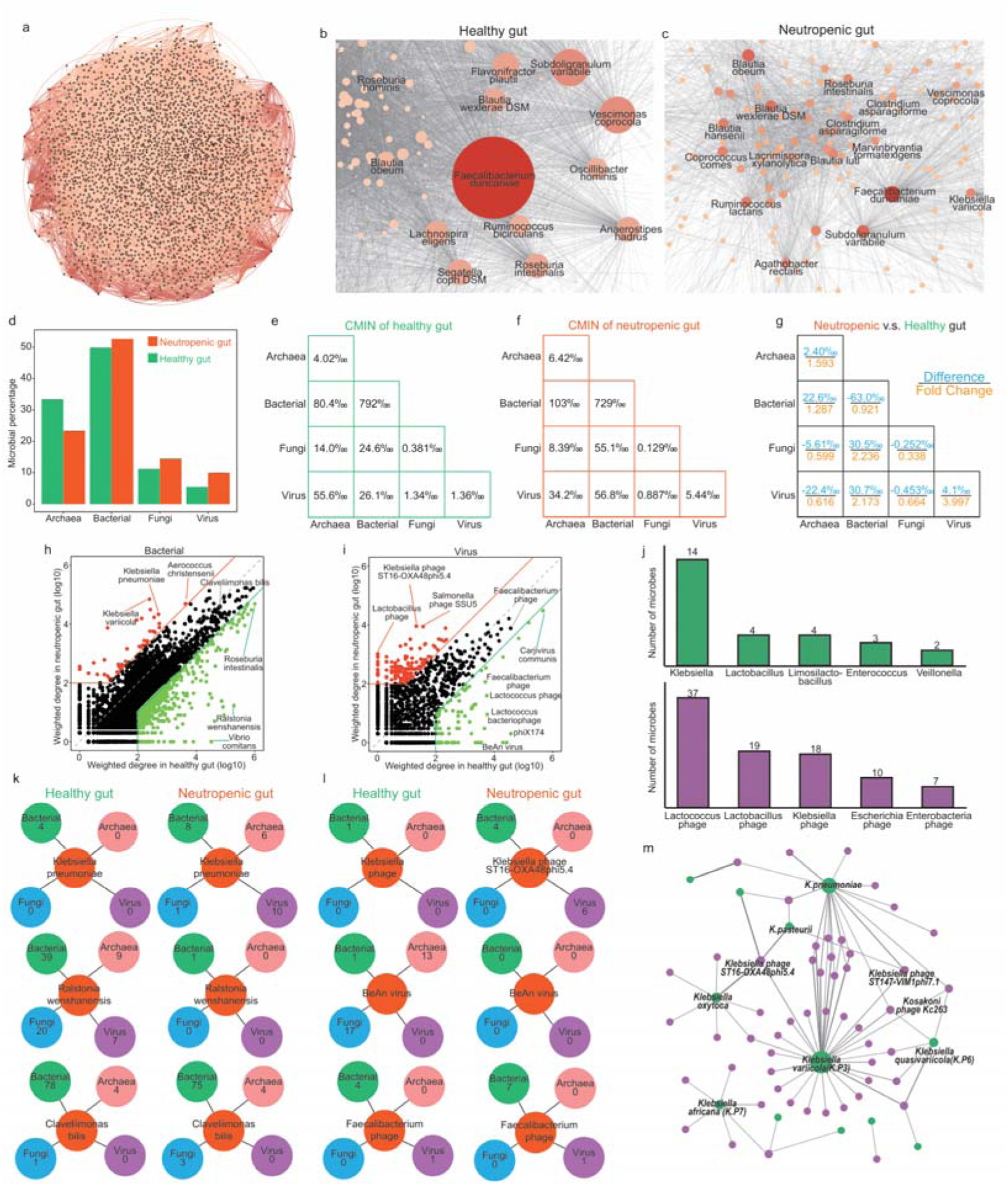
Architecture and dynamics of Cross-Domain Microbial Interaction Networks in Health and Disease. (a) The cross-domain microbial interaction network. (b,c) Force-directed network layouts comparing gut microbial interactomes in (b) healthy individuals versus (c) neutropenic patients, with node size reflecting interaction degree. (d) Stacked bar plot showing relative abundance shifts of four microbial domains between health and disease states. (e-g) Comparative analysis of interaction type proportions in healthy and neutropenic gut networks. (h, i) Differential interaction analysis between healthy and neutropenic gut networks: red indicates microbes with elevated interactions in neutropenic patients, and green indicates microbes with elevated interactions in healthy individuals. (j) Hyper-connected pathogens in neutropenic networks. (k,l) The number of interaction partners for (k) bacterial species and (l) viral strains across conditions. (m) *Klebsiella* subnetwork in neutropenic patients.

### Gut Microbial Interaction Networks in Healthy and Neutropenic Patients

To demonstrate the practical value of CMIN, we extracted gut microbial interaction subnetworks for healthy individuals and neutropenic patients (Figure 4b,c). The healthy network contained 7,825 microbial nodes connected by 22,484 interaction edges, reflecting a complex and stable ecosystem. In contrast, the neutropenic network showed marked simplification, with only 4,289 nodes and 8,133 edges. *Faecalibacterium duncanii* emerged as the key hub in both networks, yet exhibited striking topological differences between health and disease states. In healthy individuals, this species displayed high network centrality (betweenness centrality = 0.159) and maintained extensive connections with 1,223 intestinal microbes. However, in neutropenic patients, its interaction degree plummeted to 270, accompanied by reduced centrality (betweenness centrality = 0.087). Domain-level analysis revealed significant compositional shifts between groups (Figure 4d). Archaeal abundance declined substantially in neutropenic patients (health: 33.4% vs. disease: 23.4%), while bacterial populations remained stable (health: 49.9% vs. disease: 52.0%). Fungal representation increased moderately from health: 11.2% to disease: 14.5%, but the most dramatic change occurred in viral abundance, which increased by 86.7% (health: 5.4% vs. disease: 10.1%).

Cross-domain interaction analysis revealed bacterial-bacterial connections as the predominant interaction mode in both healthy and neutropenic networks, accounting for 79.2% and 72.9% of total interactions respectively, with bacterial-related interactions collectively exceeding 90% (Figure 4e-g). Viral-viral interactions exhibited the most dramatic increase (3.99-fold; health: 0.136% to disease: 0.544%), followed by viral-bacterial (2.17-fold; health: 2.61% to disease: 5.68%) and bacterial-fungal interactions (2.23-fold; health: 2.46% to disease: 5.51%). This viral interaction surge in immunocompromised states suggests enhanced viral niche expansion and potential disruption of host defense mechanisms.

Comparative analysis of bacterial and viral nodes identified distinct interaction signatures between health and disease states (Figure 4h-i). *Klebsiella spp.* emerged as hyperactive interactors in the neutropenic network, with *Klebsiella pneumoniae* showing >10-fold increased interactions. This pathogen expanded its interaction repertoire to include 6 archaeal, 8 bacterial, 1 fungal, and 10 viral partners - a dramatic increase from its limited 4 bacterial interactions in healthy network. In contrast, *Ralstonia wenshanensis* displayed healthy-network dominance, maintaining extensive archaeal (15 partners) and fungal (22 partners) connections that significantly exceeded patient-network levels. *Claveliimonas bilis* exhibited network-state stability, with interactions consistently focused within *Bacteroidetes* species (Figure 4k). Viral interaction profiling revealed *Klebsiella phage* as the third most active viral group in disease network (following *E. coli* and *Lactobacillus phages*), with *Klebsiella phage ST16-OXA48phi5.4* demonstrating broad connectivity (4 *Klebsiella* strains, 6 viral genomes). Conversely, the *BeAn virus* maintained robust interaction capacity in healthy networks (13 archaeal, 1 bacterial, 17 fungal partners), while *Faecalibacterium phage* showed network-state invariance (Figure 4l). This comparative network analysis provides insights into how host immune status reshapes microbial community architecture, revealing potential biomarkers or therapeutic targets through CMIN.

### CMIN Facilitates Phage Therapy

Phage therapy represents a promising alternative to conventional antibiotics^27^, combining targeted antimicrobial activity with low side effects and efficacy against drug-resistant strains. Leveraging our identified *Klebsiella*-phage interaction patterns, we constructed a focused subnetwork comprising 14 *Klebsiella* species and their associated phages (Figure 4m). Network analysis highlighted *Klebsiella variicola* as a superior viral interactor, establishing connections with 7 distinct phages—the broadest viral engagement among studied strains. Of particular therapeutic interest, *Klebsiella phage ST16-OXA48phi5.4* demonstrated exceptional host adaptability, interacting strongly with 4 *Klebsiella* strains. These properties position *ST16-OXA48phi5.4* as a prime candidate for combating broad-spectrum *Klebsiella* infections.

Expanding our investigation to global pathogens, we mapped interaction subnetworks for 25 WHO-certified human pathogens^28^. The *Staphylococcus aureus* subnetwork revealed three phages (StauST398-5, *phi11*, and *phiSP44-1*) exhibiting strong interactions, marking them as promising therapeutic agents for anti-staphylococcal therapy (Figure S4d). More interestingly, analysis of the *Shigella dysenteriae* subnetwork uncovered an unexpected interaction - the pathogen’s strongest phage connections involved *Enterobacteria phage fiAA91-ss* rather than traditional species-specific phages (Figure S4e). This finding challenged conventional views of phage host specificity and demonstrated how systematic cross-domain network analysis can identify unconventional phage candidates, offering new perspectives for antimicrobial treatment strategies.

### Microbial 3D Genome and Cross-domain Interactome Database and Webserver

To facilitate broad access to the generated data and enhance research reproducibility, we have curated a comprehensive dataset of microbial 3D genome architectures and cross-domain interactome. This resource is publicly accessible through a dedicated web platform (Figure S5), designed to support intuitive exploration and visualization.

The web portal provides two functionalities. First, it enables interactive browsing of the cross-domain microbial interaction network, encompassing archaea, bacteria, fungi, and viruses. Users can filter interactions by specific microbial taxa, allowing customized visualization of subnetworks relevant to their research interests (Figure 5a). Second, the platform hosts 3D structural models for 344 individual microbial genomes. These structures are presented in an interactive viewer, enabling users to explore and compare distinct 3D genome conformational types across diverse species (Figure 5b).

**Figure 5.**
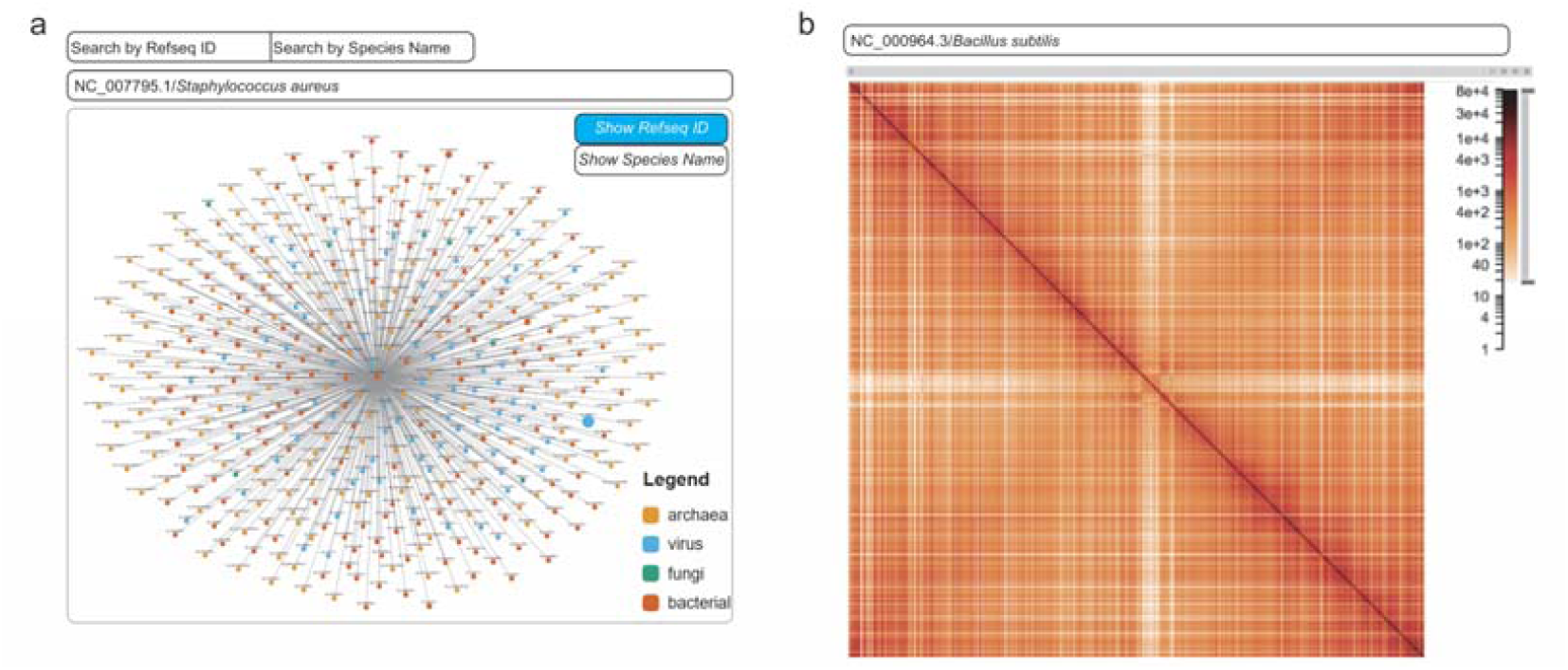
Web server for Microbe Interactome and Microbe 3D Genomes. (a) Module for exploring microbe interactome by cross-domain microbial interaction networks. (b) Module for visualizing microbe 3D genome structures using HiGlass.

## Discussion

In summary, FindMeta3D represents the first comprehensive framework capable of systematically resolving both microbial 3D genome architectures and their cross-domain interaction networks. By applying this approach to large-scale Hi-C data, we successfully reconstructed high-quality 3D genomes for 344 microbial species, revealing five distinct structural types that correlate with intrinsic sequence properties. Notably, microbial species sharing similar genome folding patterns exhibit closer phylogenetic distances, suggesting conserved mechanisms of genome organization across taxa. In human-microbe interaction analyses, FindMeta3D characterizes the contact preferences on both human and microbial genomes. Functional validation confirmed the critical role of interaction hotspots of the viral genome in mediating infection processes. The Cross-domain Microbial Interaction Network (CMIN) generated through this work offers transformative insights into microbial community dynamics, simultaneously serving as a discovery platform for pathogenic networks and a resource for identifying novel therapeutic phages.

Several limitations should be acknowledged. First, the current five classes of microbial genome folding types are constrained by the limited number of microbial species available in Hi-C data. Future studies should incorporate a more diverse array of microorganisms to enhance the generalizability and resolution of the classification system. Second, the scarcity of Hi-C data under pathological conditions restricts our understanding of disease-associated network dynamics. Deeper sequencing efforts across various disease states will be critical to elucidate these mechanisms. In our experimental validation, technical constraints necessitated the use of EBV gene-containing plasmids rather than intact viral genomes, which may not fully recapitulate native infection processes. Furthermore, while FindMeta3D assistant network analysis could identify pathogenic microbes and new therapeutic phage candidates, their clinical potential requires further evaluation.

## Conclusions

In this study, we present FindMeta3D, a powerful and scalable computational framework that enables the systematic exploration of microbial 3D genome architecture and cross-domain interaction networks from complex Hi-C datasets. By applying FindMeta3D to 528 samples, we have uncovered fundamental principles of microbial chromatin organization, identifying five evolutionarily conserved structural types linked to intrinsic genomic features. The framework further reveals species-specific preferences in microbial interactions with the human genome and identifies critical genomic hotspots essential for pathogen infectivity, as validated by functional experiments. Importantly, we construct the first Cross-domain Microbial Interaction Network (CMIN), which not only captures complex ecological and pathogenic relationships across archaea, bacteria, fungi, and viruses, but also detects dynamic network rewiring in disease states, such as enhanced Klebsiella-phage interactions in neutropenic patients. These findings highlight the regulatory significance of three-dimensional genome organization and cross-kingdom interactions in microbial communities. FindMeta3D thus provides a platform for advancing microbial genomics, understanding infection mechanisms, and guiding the development of novel antimicrobial strategies, including precision phage therapy.

## Materials and methods

### FindMeta3D algorithm

In FindMeta3D, raw Hi-C sequencing data are preprocessed using the fastp^29^ to remove low-quality reads. The parameter settings include enabling overrepresentation analysis to detect potential sequence contamination and read correction to improve alignment accuracy. Following stringent quality control, high-quality paired-end reads are aligned to the human T2T reference genome (chm13) using bwa mem^30^ or optionally bowtie2^31^, with alignment results stored in BAM format. Reads containing at least one unmapped host-aligned end are retained for subsequent microbial interaction analysis.

These unmapped reads potentially represent two interaction types: (1) host-microbe interaction (one host-mapped and one microbe-mapped end), and (2) microbial interaction (both ends mapping to microbial genomes). For systematic identification, reads undergo sequential alignment against four microbial reference databases (archaea, bacteria, fungi, and viruses) using domain-specific BWA indices. Successfully mapped reads are processed with pairtools^32^ to generate preliminary interaction files, with intra-genomic interactions used for microbial 3D structure reconstruction. Only genomes achieving ≥95% locus coverage are retained for structural analysis.

Cross-domain interactions are identified using GetCrossDomainReads.py, which extracts and merges reads mapping to different microbial domains or between host and microbial genomes. The resulting comprehensive interaction file undergoes stringent quality control, including removal of alignments with MAPQ < 10 and reads mapping to multiple species, ensuring interaction specificity by eliminating conserved regions and potential contaminants.

### Clustering of five 3D structural types

We conducted a systematic clustering analysis on the 344 identified microbial 3D genome structures. To account for potential biases caused by differences in genome length, we first standardized all genome interaction matrices to a uniform dimension of 256×256. After matrix normalization, based on the spatial distribution patterns of interactions, we manually classified the 344 3D genome structures into five structural subtypes. The classification criteria were primarily based on visual inspection of interaction heatmap patterns, including Type I (X-shape), Type II (Mammal-like), Type III (Dual X-shape), Type IV (Linear), and Type V (Circular). Then, we employed Linear Discriminant Analysis (LDA) from the scikit-learn^21^ to extract the most discriminative feature vectors that best distinguished the different types. Finally, to visualize the high-dimensional features, we applied the UMAP (Uniform Manifold Approximation and Projection) algorithm^33^ to generate a two-dimensional visualization map that revealed the distribution patterns of different microbial 3D structural types.

### Genomic bendability

we developed an algorithm called BendNet^22^, capable of calculating genomic bendability at single-nucleotide resolution. We systematically applied BendNet to all 344 microbial genomes to predict their corresponding bendability profiles. Through BendNet, we obtained high-resolution bendability maps across each genome at the single-nucleotide level.

### Identify homologs of SMC and condensin proteins in microbial genomes

We first collected all amino acid sequences of SMC and condensin proteins from the UniProt database^34^. These sequences were formatted into a standardized sequence database using makeblastdb (v2.12.0) and subsequently used to construct a BLAST alignment database^35^. Next, we employed the tblastn algorithm (E-value < 1e-5, max_target_seqs=10) to perform homology alignment between the **microbial** genome sequences and the SMC and condensin protein sequence database.

### Prediction of structural features of ACP and its variants

We extracted the alignment information of the acyl carrier protein (ACP) from the multiple sequence alignment using MAFFT^36^ and applied a majority voting strategy to determine the specific ACP variants of each structural type. Subsequently, we used the RoseTTAFold^37^ algorithm to predict the 3D structures of specific ACP variants, retaining only those with prediction errors below 0.1 Å to ensure high accuracy. Finally, the predicted structures were visualized using PyMOL, and calculate the Match Alignment Scores between pairs of structures to evaluate and compare structural similarities across different variants.

### Construction of bacterial and viral phylogenetic trees

We performed multiple sequence alignment (MSA) – MAFFT – on sequences of SMC and condensin orthologs in 301 bacterial and 41 viral genomes, respectively. Based on the MSA result, we reconstructed a maximum likelihood phylogenetic tree by IQ-TREE. During tree inference, the ModelFinder module automatically selected the best-fit evolutionary model, and 1,000 bootstrap replications were conducted to assess node support values. The final phylogenetic tree was visualized using the iTOL platform^38^.

### Calculation of microbial-human interaction frequency

To quantify the interaction frequency between microbes and human, we first counted the number of microbial-human genomic interactions for each microbial species. To eliminate potential bias caused by differences in microbial abundance, we applied a normalization strategy: the number of microbe-human interaction reads was divided by the number of intra-microbial interaction reads, yielding a normalized Interaction frequency.

### Correlation between microbial interaction preferences and epigenomic signals in human

We first identified microbial-human interaction, and then quantified the distribution of these interactions in human genome. We divided each human chromosome into bins of 1M bp. For each bin, if one microbial-human interaction was detected, the interaction preference count for that bin was increased by one. To integrate epigenetic regulatory information, the same binning strategy was applied to epigenetic marks in human cells: GM12878 (H3K36me3, H3K4me3, ATAC-seq, H3K27ac, H3K9me3), Replication Timing, nucleolar-associated domains (NADs), common fragile sites (CFSs), and lamina-associated domains (LADs). For each 1MB bin, the average signal intensity of each epigenetic mark was calculated. Finally, Pearson correlation coefficients were computed to assess the relationships between microbial interaction preferences and various epigenetic signals.

### FindMetaPeaks

We developed FindMetaPeaks to identify significant interaction hotspots based on signal processing and statistical testing. First, the microbial interaction data were represented as a one-dimensional array, where each position corresponded to a genomic coordinate and its value indicated the number of interactions at that locus. We used the find_peaks function from the SciPy library to detect local peaks in the signal. For each detected peak, we expanded outward from the summit until the signal intensity dropped to 20% of the peak height, thereby defining the boundaries of the peak region. To ensure consistent resolution across the genome, we imposed a minimum and maximum width constraint of 10 and 1000 base pairs, respectively. If a peak region did not meet these width criteria, it was either extended or shrunk accordingly. To avoid overlapping peak regions, a non-overlapping mechanism was introduced: each newly identified peak region had to start immediately after the previous one ended. Additionally, only peaks with a height exceeding a predefined standard (10 interactions) were considered for further analysis. Finally, we assessed the significance of each candidate peak using an independent t-test. The signal within the peak was compared with that in adjacent regions. Peaks with a p-value less than 0.05 were considered significant interaction hotspots and retained for downstream analysis.

### Cell Culture

Human embryonic kidney 293T (HEK293T) cells were cultured in Dulbecco’s Modified Eagle Medium (DMEM; Gibco, Cat# 11965-092) supplemented with 10% fetal bovine serum (FBS; Gibco, Cat# 10270-106), 2 mM L-glutamine (Gibco, Cat# 25032-081), and 1% penicillin-streptomycin (Gibco, Cat# 15140-122) at 37°C in a humidified incubator with 5% COD. Cells were routinely passaged every 2–3 days using 0.25% trypsin-EDTA (Gibco, Cat# 25200-072) and maintained at 70–90% confluency prior to transfection or infection experiments. Authentication of HEK293T cells was performed via short tandem repeat (STR) profiling, and cells were confirmed to be mycoplasma-negative using the MycoAlert™ Mycoplasma Detection Kit (Lonza).

### Plasmid Construction

An Epstein-Barr virus (EBV)-derived episomal vector containing the oriP replication origin and the EBNA1 gene was used to enable long-term episomal maintenance and transgene expression in mammalian cells. The vector was engineered to stably express green fluorescent protein (GFP) under the control of a constitutive promoter. Wild-type (WT) and mutant versions of the vector were generated using Gibson assembly. Mutant vectors were constructed by deleting individual interaction hotspots identified by FindMetaPeaks within the EBNA1 gene region (two hotspots) or the oriP region (three hotspots), while control vectors contained deletions in non-functional genomic regions with no predicted regulatory activity. All plasmid constructs were verified by Sanger sequencing and restriction enzyme digestion.

### Cell Transfection and Time-Course Assay

HEK293T cells were seeded in 6-well plates at a density of 2 × 10D cells per well 24 hours prior to transfection to achieve ∼70% confluency. Transfections were performed using polyethylenimine (PEI; Polysciences, Cat# 23966) at a 3:1 (μL PEI : μg DNA) ratio in serum-free Opti-MEM medium (Gibco, Cat# 31985-070). For each condition, 2 μg of plasmid DNA was transfected per well. Transfection efficiency was assessed by flow cytometry (BD FACSCanto II) at 48 hours post-transfection (day 2) to measure initial GFP expression. To evaluate long-term episomal maintenance, cells were cultured for an additional 7 days (up to day 9), with media changes every 2–3 days. GFP fluorescence intensity was monitored at day 9 to assess sustained transgene expression.

### Genomic DNA Extraction and qPCR Analysis

At day 9 post-transfection, total genomic DNA was extracted from transfected HEK293T cells using the DNeasy Blood & Tissue Kit (Qiagen, Cat# 69504) according to the manufacturer’s instructions. Quantitative real-time PCR (qPCR) was performed to measure vector copy number relative to a reference housekeeping gene GAPDH. Amplification was carried out using SYBR Green Master Mix (Applied Biosystems, Cat# 4385612). Primers specific to the GFP transgene and the reference gene were used. Absolute DNA levels were calculated using the ΔΔCt method, with results normalized to the wild-type vector control.

### RNA Isolation and RT-qPCR

Total RNA was extracted from transfected HEK293T cells at day 9 using the RNeasy Mini Kit (Qiagen, Cat# 74104) and treated with DNase I (Qiagen, Cat# 79254) to remove residual genomic DNA. Reverse transcription was performed using the High-Capacity cDNA Reverse Transcription Kit (Applied Biosystems, Cat# 4368814) with random hexamer primers. RT-qPCR was conducted using gene-specific primers for GFP and the housekeeping gene GAPDH. Gene expression levels were analyzed via the ΔΔCt method and normalized to GAPDH level.

### Flow Cytometry and Fluorescence Imaging

GFP expression was quantified by flow cytometry at both day 2 and day 9 post-transfection. A minimum of 10,000 events per sample were collected, and data were analyzed using FlowJo v10 software. Mean fluorescence intensity (MFI) was used to assess transgene expression levels. In parallel, representative images were captured using a fluorescence microscope with standardized exposure settings to visualize GFP signal across conditions.

### Construction of CMIN

After obtaining the microbial-microbial interaction data, we first counted the all interaction events between each pair of microbes. Each microbe was treated as an independent node, and the total number of interaction events between two nodes was assigned as the edge weight, thereby constructing a cross-domain microbial interaction network. To eliminate the influence of self-interaction strength on overall interaction intensity, we applied a normalization strategy: the interaction strength between any two nodes was divided by the geometric mean of their respective self-interaction strengths. This normalization removes biases caused by inherent interaction tendencies of individual nodes, ensuring comparability of interaction strengths across different pairs.

To identify statistically significant interacting events, we employed a permutation test. Specifically, we randomly shuffled the node labels of the original interaction network to generate 1,000 random networks. For each pair of nodes, we calculated the distribution of interaction strengths across these random networks and compared the observed interaction strength with this null distribution. If the p-value of the observed interaction strength fell below 0.05, the interaction pair was considered statistically significant.

## Code Available

FindMeta3D is freely available at https://github.com/JiangWenJie-stack/Findmeta3D and https://zenodo.org/records/15502445. The web server is accessible at http://wulab.bjmu.edu.cn/microbe3dgenome.

## Data available

528 Hi-C samples were obtained from the following sources: ERX2548555, ERX2548556, GSE103477, GSE113339, GSE113703, GSE116862, GSE135941, GSE137188, GSE142004, GSE145150, GSE152549, GSE160973, GSE162612, GSE163694, GSE164533, GSE189178, GSE195631, GSE198412, GSE225249, GSE225860, GSE239989, GSE98120, PRJNA1006511, PRJNA413092, PRJNA627086, PRJNA649316, PRJNA718195, GSE144475, GSE144742, GSE182881, GSE193967, GSE214856, GSE221497, GSE225771, GSE245010, GSE45966, and GSE97330.

The ChIP-seq data for histone modifications H3K36me3, H3K4me3, H3K27ac, and H3K9me3, as well as ATAC-seq data for the GM12878 cell line, were downloaded from the ENCODE database (https://www.encodeproject.org/). Replication timing data were obtained from previous studies [cite]. Nucleation-Associated Domains (NADs), Common Fragile Sites (CFS), and Lamina-Associated Domains (LADs) were retrieved from the UCSC Genome Browser.

## Acknowledgment

This work was supported by the National Natural Science Foundation of China (32270683 and 32470662); the Beijing Natural Science Foundation (5242006); the Fundamental Research Funds for the Central Universities (BMU2021YJ064) to H.J.W.; CAMS Innovation Fund for Medical Sciences (2021-I2M-5-003) to M.X.; the Science Foundation of Peking University Cancer Hospital (ZY202418) to Y.C.S.; the China Postdoctoral Science Foundation (2024M750125) to N.N.W.; We gratefully acknowledge the High-performance Computing Platform of Peking University for conducting the data analyses.

## Reference

1. Jerkovic, I., and Cavalli, G. (2021). Understanding 3D genome organization by multidisciplinary methods. Nat Rev Mol Cell Biol 22, 511–528. 10.1038/s41580-021-00362-w.

2. Xu, J., Song, F., Lyu, H., Kobayashi, M., Zhang, B., Zhao, Z., Hou, Y., Wang, X., Luan, Y., Jia, B., et al. (2022). Subtype-specific 3D genome alteration in acute myeloid leukaemia. Nature 611, 387–398. 10.1038/s41586-022-05365-x.

3. Yang, H., Luan, Y., Liu, T., Lee, H.J., Fang, L., Wang, Y., Wang, X., Zhang, B., Jin, Q., Ang, K.C., et al. (2020). A map of cis-regulatory elements and 3D genome structures in zebrafish. Nature 588, 337–343. 10.1038/s41586-020-2962-9.

4. Ona, L., Shreekar, S.K., and Kost, C. (2025). Disentangling microbial interaction networks. Trends Microbiol. 10.1016/j.tim.2025.01.013.

5. Cao, C., Yue, S., Lu, A., and Liang, C. (2024). Host-Gut Microbiota Metabolic Interactions and Their Role in Precision Diagnosis and Treatment of Gastrointestinal Cancers. Pharmacol Res 207, 107321. 10.1016/j.phrs.2024.107321.

6. Wang, S., Mu, L., Yu, C., He, Y., Hu, X., Jiao, Y., Xu, Z., You, S., Liu, S.L., and Bao, H. (2024). Microbial collaborations and conflicts: unraveling interactions in the gut ecosystem. Gut Microbes 16, 2296603. 10.1080/19490976.2023.2296603.

7. Brito, I.L. (2021). Examining horizontal gene transfer in microbial communities. Nat Rev Microbiol 19, 442–453. 10.1038/s41579-021-00534-7.

8. Gao, S., Khan, M.I., Kalsoom, F., Liu, Z., Chen, Y., and Chen, Z. (2022). Role of gene regulation and inter species interaction as a key factor in gut microbiota adaptation. Arch Microbiol 204, 342. 10.1007/s00203-022-02935-5.

9. Sulaiman, J.E., Thompson, J., Cheung, P.L.K., Qian, Y., Mill, J., James, I., Im, H., Vivas, E.I., Simcox, J., and Venturelli, O.S. (2025). Phocaeicola vulgatus shapes the long-term growth dynamics and evolutionary adaptations of Clostridioides difficile. Cell Host Microbe 33, 42–58 e10. 10.1016/j.chom.2024.12.001.

10. Zomorrodi, A.R., and Segre, D. (2017). Genome-driven evolutionary game theory helps understand the rise of metabolic interdependencies in microbial communities. Nat Commun 8, 1563. 10.1038/s41467-017-01407-5.

11. Xu, Y., Yin, H., Jiang, H., Liang, Y., Guo, X., Ma, L., Xiao, Y., and Liu, X. (2013). Comparative study of nickel resistance of pure culture and co-culture of Acidithiobacillus thiooxidans and Leptospirillum ferriphilum. Arch Microbiol 195, 637–646. 10.1007/s00203-013-0900-z.

12. Le, H.H., Lee, M.T., Besler, K.R., Comrie, J.M.C., and Johnson, E.L. (2022). Characterization of interactions of dietary cholesterol with the murine and human gut microbiome. Nat Microbiol 7, 1390–1403. 10.1038/s41564-022-01195-9.

13. Soucy, S.M., Huang, J., and Gogarten, J.P. (2015). Horizontal gene transfer: building the web of life. Nat Rev Genet 16, 472–482. 10.1038/nrg3962.

14. Moura de Sousa, J., Lourenco, M., and Gordo, I. (2023). Horizontal gene transfer among host-associated microbes. Cell Host Microbe 31, 513–527. 10.1016/j.chom.2023.03.017.

15. Lieberman-Aiden, E., van Berkum, N.L., Williams, L., Imakaev, M., Ragoczy, T., Telling, A., Amit, I., Lajoie, B.R., Sabo, P.J., Dorschner, M.O., et al. (2009). Comprehensive mapping of long-range interactions reveals folding principles of the human genome. Science 326, 289–293. 10.1126/science.1181369.

16. Marbouty, M., Thierry, A., Millot, G.A., and Koszul, R. (2021). MetaHiC phage-bacteria infection network reveals active cycling phages of the healthy human gut. Elife 10. 10.7554/eLife.60608.

17. Philippot, L., Chenu, C., Kappler, A., Rillig, M.C., and Fierer, N. (2024). The interplay between microbial communities and soil properties. Nat Rev Microbiol 22, 226–239. 10.1038/s41579-023-00980-5.

18. Wu, R., Davison, M.R., Nelson, W.C., Smith, M.L., Lipton, M.S., Jansson, J.K., McClure, R.S., McDermott, J.E., and Hofmockel, K.S. (2023). Hi-C metagenome sequencing reveals soil phage-host interactions. Nat Commun 14, 7666. 10.1038/s41467-023-42967-z.

19. Nurk, S., Koren, S., Rhie, A., Rautiainen, M., Bzikadze, A.V., Mikheenko, A., Vollger, M.R., Altemose, N., Uralsky, L., Gershman, A., et al. (2022). The complete sequence of a human genome. Science 376, 44–53. 10.1126/science.abj6987.

20. Jiang, W.-J., Cai, K., Sun, Y., Liu, A., Zhu, H., Gao, R., Zhong, C., Wei, N., Lai, F., Fei, T., et al. (2025). Harmonizing single cell 3D genome data with STARK and scNucleome. bioRxiv, 2025.2005.2010.653247. 10.1101/2025.05.10.653247.

21. Pedregosa, F., Varoquaux, G., Gramfort, A., Michel, V., Thirion, B., Grisel, O., Blondel, M., Prettenhofer, P., Weiss, R., Dubourg, V., et al. (2011). Scikit-learn: Machine Learning in Python. J. Mach. Learn. Res. 12, 2825–2830.

22. Jiang, W.J., Hu, C., Lai, F., Pang, W., Yi, X., Xu, Q., Wang, H., Zhou, J., Zhu, H., Zhong, C., et al. (2023). Assessing base-resolution DNA mechanics on the genome scale. Nucleic Acids Res 51, 9552–9566. 10.1093/nar/gkad720.

23. Lioy, V.S., Cournac, A., Marbouty, M., Duigou, S., Mozziconacci, J., Espeli, O., Boccard, F., and Koszul, R. (2018). Multiscale Structuring of the E. coli Chromosome by Nucleoid-Associated and Condensin Proteins. Cell 172, 771–783 e718. 10.1016/j.cell.2017.12.027.

24. Hirano, T. (2016). Condensin-Based Chromosome Organization from Bacteria to Vertebrates. Cell 164, 847–857. 10.1016/j.cell.2016.01.033.

25. Roberts, D.M., Anchimiuk, A., Kloosterman, T.G., Murray, H., Wu, L.J., Gruber, S., and Errington, J. (2022). Chromosome remodelling by SMC/Condensin in B. subtilis is regulated by monomeric Soj/ParA during growth and sporulation. Proc Natl Acad Sci U S A 119, e2204042119. 10.1073/pnas.2204042119.

26. Minh, B.Q., Schmidt, H.A., Chernomor, O., Schrempf, D., Woodhams, M.D., von Haeseler, A., and Lanfear, R. (2020). IQ-TREE 2: New Models and Efficient Methods for Phylogenetic Inference in the Genomic Era. Mol Biol Evol 37, 1530–1534. 10.1093/molbev/msaa015.

27. MacNair, C.R., Rutherford, S.T., and Tan, M.W. (2024). Alternative therapeutic strategies to treat antibiotic-resistant pathogens. Nat Rev Microbiol 22, 262–275. 10.1038/s41579-023-00993-0.

28. Jesudason, T. (2024). WHO publishes updated list of bacterial priority pathogens. Lancet Microbe 5, 100940. 10.1016/j.lanmic.2024.07.003.

29. Chen, S. (2023). Ultrafast one-pass FASTQ data preprocessing, quality control, and deduplication using fastp. Imeta 2, e107. 10.1002/imt2.107.

30. Li, H., and Durbin, R. (2010). Fast and accurate long-read alignment with Burrows-Wheeler transform. Bioinformatics 26, 589–595. 10.1093/bioinformatics/btp698.

31. Langmead, B., Wilks, C., Antonescu, V., and Charles, R. (2019). Scaling read aligners to hundreds of threads on general-purpose processors. Bioinformatics 35, 421–432. 10.1093/bioinformatics/bty648.

32. Open2C, Abdennur, N., Fudenberg, G., Flyamer, I.M., Galitsyna, A.A., Goloborodko, A., Imakaev, M., and Venev, S.V. (2024). Pairtools: From sequencing data to chromosome contacts. PLoS Comput Biol 20, e1012164. 10.1371/journal.pcbi.1012164.

33. Narayan, A., Berger, B., and Cho, H. (2021). Assessing single-cell transcriptomic variability through density-preserving data visualization. Nat Biotechnol 39, 765–774. 10.1038/s41587-020-00801-7.

34. UniProt, C. (2025). UniProt: the Universal Protein Knowledgebase in 2025. Nucleic Acids Res 53, D609–D617. 10.1093/nar/gkae1010.

35. Camacho, C., Coulouris, G., Avagyan, V., Ma, N., Papadopoulos, J., Bealer, K., and Madden, T.L. (2009). BLAST+: architecture and applications. BMC Bioinformatics 10, 421. 10.1186/1471-2105-10-421.

36. Katoh, K., Rozewicki, J., and Yamada, K.D. (2019). MAFFT online service: multiple sequence alignment, interactive sequence choice and visualization. Brief Bioinform 20, 1160–1166. 10.1093/bib/bbx108.

37. Baek, M., DiMaio, F., Anishchenko, I., Dauparas, J., Ovchinnikov, S., Lee, G.R., Wang, J., Cong, Q., Kinch, L.N., Schaeffer, R.D., et al. (2021). Accurate prediction of protein structures and interactions using a three-track neural network. Science 373, 871–876. 10.1126/science.abj8754.

38. Letunic, I., and Bork, P. (2024). Interactive Tree of Life (iTOL) v6: recent updates to the phylogenetic tree display and annotation tool. Nucleic Acids Res 52, W78–W82. 10.1093/nar/gkae268.

